# The reality of “food porn”: Larger brain responses to food-related cues than to erotic images predict cue-induced eating

**DOI:** 10.1101/184838

**Authors:** Francesco Versace, David W. Frank, Elise M. Stevens, Menton M. Deweese, Michele Guindani, Susan M. Schembre

## Abstract

While some individuals can defy the lure of temptation, many others find appetizing food irresistible. Using event-related potentials, we showed that individuals who find food-related images more motivationally relevant than erotic ones (“sign-trackers”) are more susceptible to cue-induced eating and, in the presence of a palatable food option, eat twice as much as individuals with the opposite brain reactivity profile (“goal-trackers”). These findings contribute to the understanding of the neurobiological basis of vulnerability to cue-induced behaviors.

---

Over 62,000 photos are shared worldwide each day under #foodporn^1^. These images glamorize the high-calorie, highly palatable foods that are believed to encourage the maladaptive eating patterns that contribute to today’s obesity epidemic. Researchers posit that food-related stimuli elicit compulsive eating, thereby promoting excessive energy intake^2^. Yet, attempts to test this hypothesis yielded to inconsistent findings^3^, suggesting that the neural underpinnings of human cue-induced maladaptive behaviors remain unclear.

Individual differences in susceptibility to cue-induced behaviors can explain part of these inconsistencies. Preclinical models show that rats prone to attributing incentive salience to discrete food-related cues (“sign-trackers”) are more vulnerable to compulsive cue-induced behaviors than rats not prone to do so (“goal-trackers”)^4^. Incentive salience refers to the motivational properties that grant stimuli the power to capture attention, activate affective states, and motivate behaviors^5^. We hypothesized that, similar to animal models, humans who attribute more incentive salience to food-related cues than to erotic images would be more susceptible to cue-induced eating. To test this hypothesis, we measured brain responses and eating behavior during a cued food delivery task^6^ in 49 volunteers from the community (aged 24 to 65 years, 45% female; 41% overweight, 37% obese). During the task (see **Supplementary Methods** and **Supplementary Figure 1**), we collected event-related potentials (ERPs) while participants viewed a slideshow that included images depicting high and low arousing pleasant (erotica and romantic), neutral (household objects and individuals engaged in mundane activities), high and low arousing unpleasant (mutilations, violence, pollution), and highly-palatable food-related contents (high-calorie sweet and savory food). We told participants that the slideshow included images of sweet and savory foods and that after each image of a sweet food, a machine would dispense one chocolate candy (Food-paired trials); whereas, no candies would follow the images showing savory foods (Food-unpaired trials). The food category/candy delivery pairing was counterbalanced across subjects. Participants decided whether to eat or discard each of the 60 candies delivered during the task.

**Figure 1:**
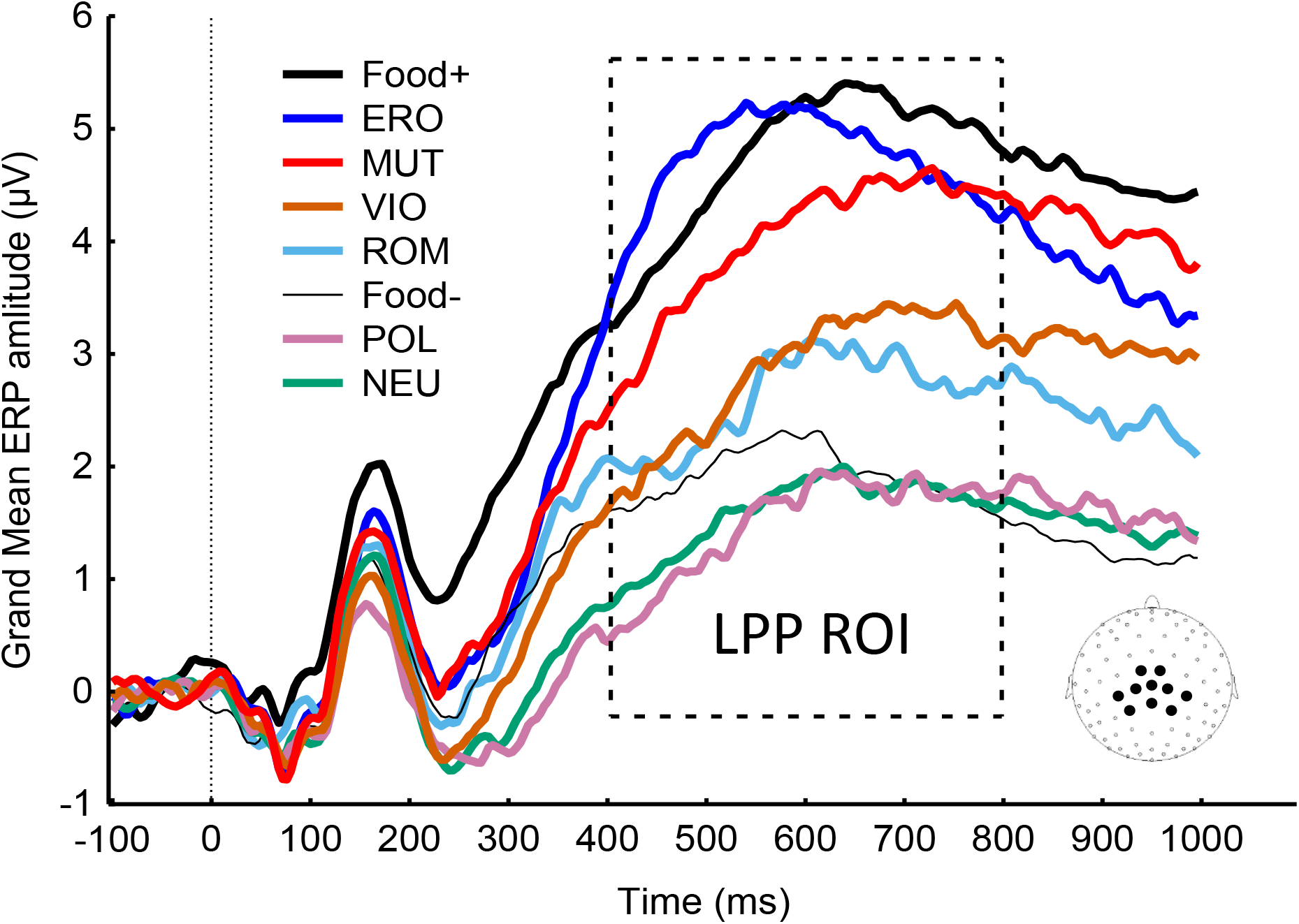
ERPs from centro-parietal sites (see inset for electrode location) show that, on average, both pleasant and unpleasant highly-relevant contents increase the amplitude of the Late Positive Potential (LPP). The box shows the time region of interest (ROI) used to compute the LPP amplitude. Note: Food+: food images paired with food delivery, Food-: food images not paired with food delivery, ERO: erotica, ROM: romance, NEU: neutral, POL: pollution, VIO: violence, MUT: mutilations.

First, we classified participants using their brain responses to the image categories (see also **Supplementary Methods**). Then, this classification was used to predict the rate at which participants ate the candies. For each participant, we calculated the amplitude of the late positive potential (LPP) evoked by each image category (i.e., Food-paired, Erotica, Romantic, Food-unpaired, Neutral, Pollution, Violence, Mutilations; **Figure 1**). The LPP is a sustained ERP component that peaks between 400 and 800 ms over central and parietal sites and is considered the most replicable and reliable electrophysiological index of motivational relevance^7^. As expected, the mean amplitude of the LPP increased as a function of the images’ motivational relevance (LPP to erotica and mutilations > LPP to romantic and violence > LPP to neutral contents; all Ps<.0001). We classified participants by applying *k*-means cluster analysis to their LPP responses. Cluster analysis is an unsupervised multivariate technique used to classify objects (i.e., participants) based on their characteristics (i.e., the participants’ brain responses)^8^. The optimal solution (see **Supplementary Results**) identified two groups that we labeled as “sign-trackers” and “goal-trackers”. **Figure 2 (left panel)** shows that sign-trackers (N=20, 41%) had larger LPPs to food-paired images than to erotic images (P<.0001); whereas, goal-trackers (N=29, 59%) had larger LPPs to erotic images than to food-paired images (P<.0001). Both groups showed the typical affective reactivity pattern such that, irrespective of hedonic content, the amplitude of the LPP increased as a function of the images’ motivational relevance. Excluding food images, the quadratic trend was significant for both sign - and goal-trackers (Ps<.0001) and the two groups had comparable LPP responses to all categories of stimuli except to food-paired images (P<.0001). These results indicate that, although every participant was aware that food-paired stimuli predicted food delivery, only sign-trackers attributed more motivational relevance to food-paired images than to erotic images. Conversely, goal-trackers processed food-paired stimuli as though they had low motivational relevance. Finally, **Figure 2 (right panel)** shows that sign-trackers ate more than twice as many candies as goal-trackers (21 vs. 8; U=188.5, P=.036). In the quasi-Poisson linear generalized regression model, sign-trackers ate candies at a rate that was 2.2 times greater (95% CI: 1.14, 4.37; P=.024) than that of goal-trackers, after adjusting for potential confounders (age, BMI, gender, and pre-experiment hunger level, see **Supplementary Results**).

**Figure 2:**
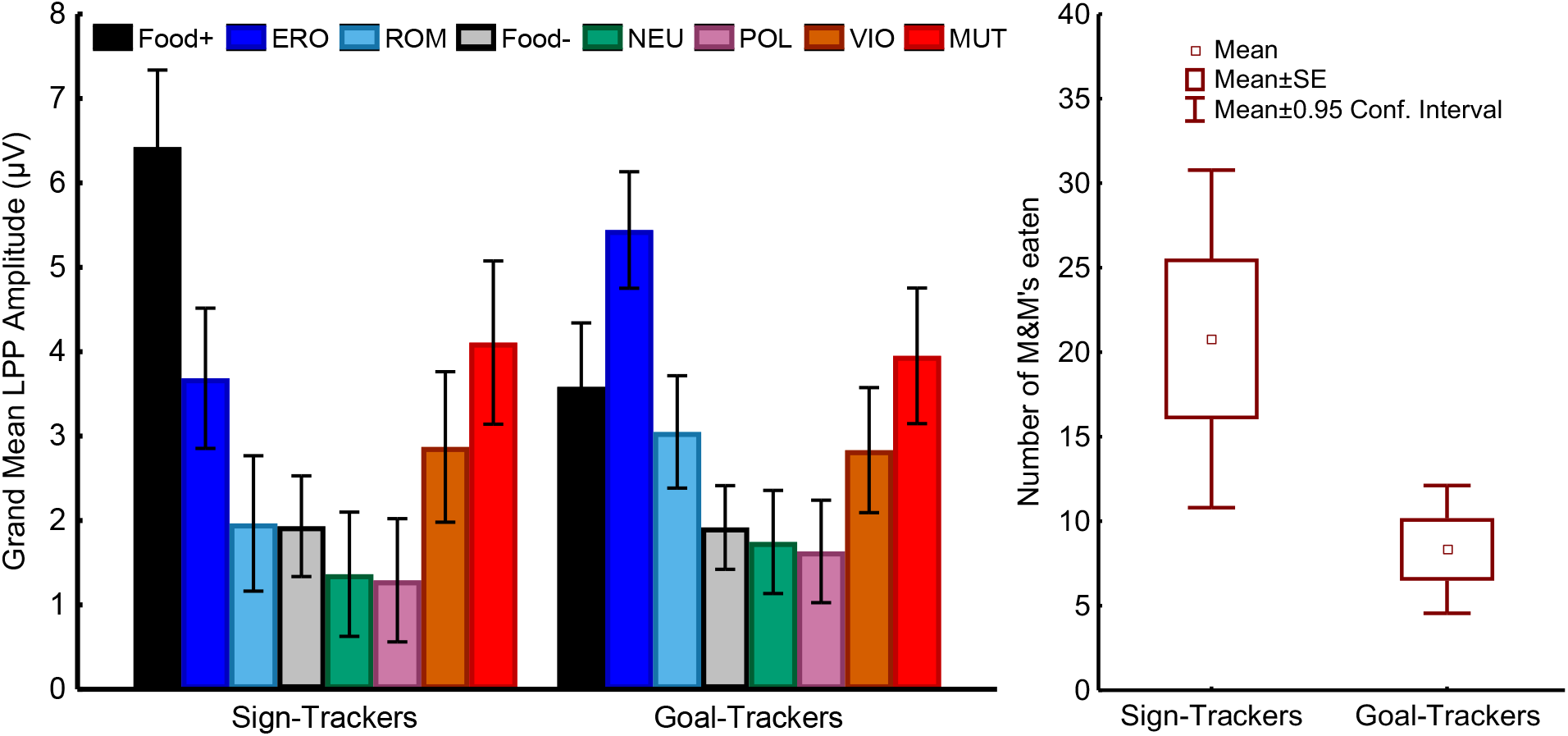
**Left:** The *k*-means cluster analysis performed on the LPP responses yielded two clusters fitting with the hypothesized sign-vs. goal-tracking dichotomy. Sign-trackers (N=20) attributed more motivational relevance to Food+ images than to erotic images while Goal-trackers (N=29) had the opposite brain reactivity pattern. **Right:** Sign-trackers ate more than twice as many candies as goal-trackers (U=188.5, P=.036). Note: Food+: food images paired with food delivery, Food-: food images not paired with food delivery, ERO: erotica, ROM: romance, NEU: neutral, POL: pollution, VIO: violence, MUT: mutilations; LPP: Late Positive Potential.

By directly measuring brain responses to a wide array of motivationally relevant stimuli, we showed that individuals who attribute more motivational relevance to food-related than to erotic images are more susceptible to cue-induced eating. This neurobehavioral pattern mirrors what is observed in animal models during Pavlovian conditioning, where sign-tracking behavior is associated with larger phasic dopamine responses to stimuli predicting rewards than to actual rewards in the nucleus accumbens^9^. While several factors motivate food consumption, these data show that attributing high motivational relevance to food-related cues significantly increases the likelihood of an individual engaging in maladaptive eating behaviors, especially when this trait co-occurs with high impulsivity or with a weakened ability to regulate reactions to cues. By contributing to the understanding of the biological basis underlying individual differences in vulnerability to cue-induced eating, these findings represent a step toward identifying new targets for personalized weight control interventions aimed at regulating the intense motivation to eat that some individuals experience in the presence of highly palatable foods.

## ACKNOWLEDGMENTS

This work was supported by the National Institute on Drug Abuse award R01-DA032581 to FV, by MD Anderson funds to FV and SMS (CA-016672). MMD was supported by the National Cancer Institute of the National Institutes of Health under Award Number R25CA057730. The content is solely the responsibility of the authors and does not necessarily represent the official views of the NIH. We thank Kimberly Claiborne, Jennifer Ng, and Danika Dirba for their help during data collection.

## Supplementary Methods

### Participants

We recruited 60 participants from the Houston metropolitan area using flyers, magazine, and newspaper advertisements. Participants were eligible for the study if they were between 18 and 65 years of age, were neither pregnant nor breast feeding, did not report any history of psychiatric disorders, seizures, head injuries with loss of consciousness, uncorrected visual impairments, eating disorders, allergies, or diet-related chronic diseases that might have prevented consumption of chocolate candies. All participants received monetary compensation for their time and for parking/travel, totaling $60. Due to poor recording quality, largely attributed to excessive movement during the task, 11 participants were excluded at various stages of the data reduction procedures, leaving 49 participants in the final sample.

### Procedures

The study included a telephone interview to verify study eligibility, followed by one in-person laboratory visit. At the in-person visit, a research assistant reviewed the study with the participant and obtained informed consent. Then, the research assistant measured the participant’s weight and height, and, using a computer-assisted procedure, collected answers to a series of questionnaires about impulsivity, eating behaviors, hunger, mood, and hedonic capacity. At the completion of the questionnaire assessment, the research assistant placed the sensors for the electroencephalographic (EEG) recording and described the task to the participant. To ensure that the participant fully understood the instructions, the research assistant remained in the room while 11 test trials were carried out (2 candies were delivered during this phase), then the research assistant left the room and began the EEG recording. At the end of the EEG session, the research assistant removed the sensors, debriefed and compensated the participant. All study procedures were approved by the University of Texas MD Anderson Cancer Center Institutional Review Board.

### Questionnaires

The computerized battery included the following questionnaires:

*Barratt Impulsiveness Scale* The Barratt Impulsiveness Scale^1^ (BIS) includes 30 items describing common impulsive or non-impulsive behaviors and preferences designed to assess the personality/behavioral construct of impulsiveness.

*Weight-Related Eating Questionnaire.* The 16-item Weight-Related Eating Questionnaire^2^ (WREQ) assesses four theory-based aspects of eating behavior labeled compensatory restraint, routine restraint, susceptibility to external cues, and emotional eating.

*Center for Epidemiological Studies Depression Scale (brief).* The Center for Epidemiological Studies Depression Scale^3^ (brief) is a 10-item self-report instrument assessing the frequency of several depressive symptoms and originally developed for studying depressive symptomatology in the general population.

*Positive and Negative Affect Schedule (PANAS).* The PANAS^4^ is a 20-item self-report instrument designed to measure the two primary measures of mood: positive and negative affect. This instrument is a reliable and valid measure of the two mood constructs ^5^.

*Snaith-Hamilton Pleasure Scale (SHAPS).* The SHAPS^6^ is a self-report measure of anhedonia that, unlike other instruments, was specifically developed to be unaffected by social class, gender, age, dietary habits, or nationality. The SHAPS is a reliable and valid questionnaire to assess hedonic tone in patient and non-patient populations ^7^ *Satiety Labeled Intensity Magnitude (SLIM)^8^* is a sensitive, reliable, and easy to-use scale for measuring perceived satiety.

### Cued food delivery task

During the cued food delivery task, participants viewed a series of images presented with a computer using E-Prime software (version 2.0.8.74; PST Inc., Pittsburgh, PA) on a 17-inch LCD monitor. M&M’s^®^ chocolate candies were delivered in a receptacle within arm’s reach from the participant, situated to the right of the computer monitor^9^ (**Figure S1**). Picture stimuli consisted of eight categories covering a variety of content: neutral (people, objects), highly arousing pleasant (erotica) and unpleasant (mutilations), low arousing pleasant (romantic) and unpleasant (violence), unpleasant objects (pollution), and palatable food (sweet, savory). The images were selected from the International Affective Picture System^10^ and from a database of images that we used in previous studies from our laboratory^11^. The task was divided into 6 equivalent blocks lasting approximately 5 minutes each. Each block included the pseudo-random (i.e., no more than two consecutive images of the same category) presentation of 55 images: 10 neutral, 10 pleasant (5 erotica and 5 romantic), 15 unpleasant (5 mutilations, 5 violence, 5 pollution), and 20 food-related (10 sweet and 10 savory). Images were not repeated during the task. One category of food images (either sweet or savory, counterbalanced across subjects) was designated as the “food-paired” image: 1000 ms after image onset, an M&M’s^®^ chocolate candy was released from a dispenser and, through a tube, was delivered in a receptacle where the participant could pick it up and either eat it or deposit it in a box. The food predictive image remained visible on the screen until the participant either pushed a button to indicate having swallowed the candy or until the candy was deposited in the deposit box. All other images, including the “food unpaired” images (i.e., images of food not followed by candy delivery), were presented for 2200 ms. A random inter-trial interval of 500-2000 ms separated each trial. **Figure S2** illustrates the sequence of events during the task. Instructions at the beginning of the task indicated to the participant which food category was designated as the food predictive category. In this way, the participant did not have to learn the contingency and all trials could be used in the analyses. During the test trials that preceded the task, we presented 11 images and two candies were delivered.

### EEG acquisition and data reduction

During the task, we recorded EEG continuously using a 129-channel Geodesic Sensor Net, amplified with an AC-coupled high input impedance (200 MΩ) amplifier (Geodesic EEG System 200; Electrical Geodesics Inc., Eugene, OR), and referenced to Cz. The sampling rate was 250 Hz, and data were filtered online by using 0.1 Hz high-pass and 100 Hz low-pass filters. Scalp impedance of each sensor was kept below 50 KΩ, as suggested by the manufacturer.

After EEG data collection, we filtered the data with a 30-Hz low-pass filter, inspected the EEG traces to evaluate the quality of the recording, and identified and interpolated (using spherical splines) channels contaminated by artifacts for more than 50% of the recording time. At this stage, we discarded three participants due poor EEG quality of the recording. For the remaining 57 participants, we corrected eye blinks using a spatial filtering method as implemented in BESA ver. 5.1.8.10 (MEGIS Software GmbH, Gräfelfing, Germany). After eye blink correction, we transformed the EEG data to the average reference and segmented the data using the picture onset to time-lock the ERPs. The segments started 100 ms before picture onset and ended 1100 ms later. Baseline was defined as the 100 ms interval preceding picture onset. Artifacts affecting sensors within each trial were identified using the following criteria: EEG amplitude above 100 or below-100 μV; absolute voltage difference between any two data points within the segment larger than 100 μV; voltage difference between two contiguous data points above 25 μV; and less than 0.5 μV variation for more than 100 ms. A segment was excluded from the subsequent averages if more than 10% of the sensors within the segment were contaminated by artifacts. At the end of this process, the average ERPs were calculated at each scalp site for each picture category. If a participant had fewer than 20% of the possible trials included in any category average, the participant was excluded from the subsequent analyses (eight participants were excluded at this stage, leaving 49 participants in the sample). We used the amplitude of the late positive potential (LPP) as a measure of motivational relevance^12,13^. The LPP for each picture category for each participant was calculated by averaging the voltage recorded between 400 and 800 ms after picture onset from 10 central and parietal sensors (EGI HydroCel Geodesic Sensor Net sensors: 7, 31, 37, 54, 55, 79, 80, 87, 106, 129). This group of sensors, the same that we used in our previous studies^11,14,15^, covers the area where the LPP amplitude differences between neutral and emotional pictures is maximal. Additionally, a preliminary analysis showed that the amplitude of the LPPs for neutral stimuli depicting objects or people was comparable; hence, we decided to collapse the two neutral categories together.

### Classification of participants as sign-trackers or goal-trackers

To classify participants as sign-or goal-trackers, we used the same procedure that we followed in our previous studies^11,15,16^. For each individual, we calculated the mean LPP evoked by each stimulus category (i.e., food paired, erotica, romantic, food unpaired, neutral, disgust, violence, and mutilations) between 400 and 800 ms over 10 centroparietal sensors. To account for individual variation in absolute voltage amplitude we standardized the LPP values using ipsatization^17^. Then, we classified individuals based on their brain reactivity profiles using *k*-means cluster analysis as implemented in the R statistical package^18^. Cluster analysis is a multivariate technique that groups individuals by minimizing within-groups variability and maximizing between-groups variability^19^. The algorithm is unsupervised, using as constraints only the number of clusters and the variables used for deriving the solution. The optimal number of cluster and corresponding classification was assessed using the Silhouette coefficient method^20^ and the gap statistics^21^. The groups extracted using cluster analysis can differ with respect to any brain reactivity pattern, hence the first analytic steps consisted of a series of post-analysis validation checks (see below) aimed at confirming **a)** the reliability of using the amplitude of the LPP to measure the motivational relevance of the visual stimuli used in the experiment and **b)** the replicability of the sign-vs. goal-tracking categorization based on LPP responses.

## Statistical Analyses

### Event-Related Potentials

The first post-analysis validation check tested whether both groups extracted using k-means cluster analysis showed increasingly larger LPPs for images with increasing motivational relevance (i.e., erotica and mutilations > romantic and violence > neutral and pollution). Within each group, we tested the presence of a quadratic trend using polynomial contrasts. The second validation check tested whether the two brain reactivity profiles extracted using cluster analysis fit the sign-vs. goal-tracking dichotomy (i.e., one group shows higher reactivity to food-predictive images than to pleasant images, and the other group shows the opposite pattern) and whether the two groups differed in reactivity to any image category. To conduct these tests, we ran an analysis of variance (ANOVA) using the amplitude of the LPP as the dependent variable, the eight picture categories (food-predictive, erotica, romantic, food non-predictive, neutral, pollution, violence, mutilations) as a within subjects factor, and the two groups (goal-trackers vs. sign-trackers) as a between subjects factor. To account for a violation of sphericity, we used multivariate ANOVA^22^. We used pairwise comparisons with Bonferroni correction to test for the presence of significant differences among categories within and between groups.

### Self-report questionnaires and demographics

To test whether sign-trackers and goal-trackers differed in age, gender, BMI, impulsivity, and mood, we conducted ANOVAs on these variables. The level of satiety before and after the session was compared between the two groups using ANOVA and, for each group, we tested whether there was a significant difference from the “neither hungry nor full” anchor point before and after the session.

### Eating behavior

To test for the presence of statistically significant differences between sign-trackers and goal-trackers in the number of candies eaten by each participant during the experiment, we used the nonparametric Mann-Whitney U test. Then, to take into account over-dispersion in the data, we also assessed the statistical significance of the differences in eating behavior in the two clusters using a quasi-Poisson generalized linear regression model with a scale dispersion parameter. Finally, we adjusted the analysis for the influence of potential confounding variables on eating behavior, by considering age, gender, BMI, and level of appetite pre-experiment as additional covariates in the Poisson generalized linear regression model.

## Supplementary Results

### Demographics and Self-report Questionnaires

**Supplementary Table 1** shows demographic and self-report variables separated by sign-and goal-trackers. The two groups did not differ in terms of gender distribution (P=.18), age (P=.35), or BMI (P=.74). On the Barratt Impulsivity scale (BIS), the total score suggested that sign-trackers were somewhat more impulsive than goal-trackers, but the differences were not statistically significant (P=.15). The analyses conducted on the BIS subscales showed that sign-trackers were significantly (P<.05) more impulsive than goal-trackers in the attentional impulsiveness subscale (i.e., “focusing on the task at hand” and “thought insertion and racing thoughts”) and the non-planning impulsiveness subscale (i.e., “planning and thinking carefully” and “enjoying challenging mental tasks”), but not on the motor impulsiveness subscale (i.e., “acting on the spur of the moment” and “a consistent life style”). Mood questionnaires (CES-D, PANAS, SHAPS) did not show any significant difference between groups (all Ps>.18). The WREQ total score and the scores to its subscales were similar in the two groups (all Ps>.27). The level of satiety in the two groups was similar before (P>.44) and after the session (P>.49).

### Cluster analysis

**Supplementary Figures 3 and 4** show the results of the silhouette method and the gap statistic method to choose the optimal number of clusters on the k-means clustering algorithm, as implemented in the R module “factoextra”^23^. Both criteria indicated that the two-cluster solution is the most appropriate.

### Analysis of eating behavior patterns and brain reactivity

**Supplementary Table 2** shows the results of an unadjusted quasi Poisson generalized linear regression model (quasi-Poisson GLM) relating the number of candies eaten by the participants to their cluster assignment. The quasi Poisson GLM relies on a log link to relate the regression equation to the count response, by positing *E(Y* |X) = *μ,log(μ*) = *X'β.* The coefficients of the regression identify changes in the response rate for a unit-increase in the corresponding covariate on the log-scale, with respect to the reference baseline. **Table 3** shows the results of the quasi-Poisson GLM when adjusting for known potential confounders (age, gender, BMI and level of appetite at the beginning of the experiment).

**Supplementary Table 1.** Participant demographic information and questionnaire scores by cluster membership

| Characteristic | All<br>(n=49) | Goal-trackers<br>(n=29) | Sign-trackers<br>(n=20) | P-value |
| --- | --- | --- | --- | --- |
| Age (years) | 47 | 48 | 46 | *P*=.46 |
| Women | 45% | 51% | 35% | *P*=.25 |
| Race |  |  |  |  |
| African American | 67% | 62% | 75% |  |
| Caucasian | 26% | 31% | 20% |  |
| Other | 7% | 7% | 5% |  |
| BMI | 31 | 31 | 31 |  |
| BIS |  |  |  |  |
| Attentional | 14.16 | 13.07 | 15.75 | *P*=.01 |
| Motor | 21.49 | 21.14 | 22.00 | *P*=.49 |
| Non planning | 23.06 | 21.62 | 25.15 | *P*=.04 |
| CESD | 7.20 | 6.48 | 8.25 | *P*=.24 |
| SHAPS | 47.55 | 47.14 | 48.15 | *P*=.53 |
| PANAS + | 34.63 | 34.14 | 35.35 | *P*=.65 |
| PANAS - | 17.39 | 16.21 | 19.10 | *P*=.18 |
| WREQ |  |  |  |  |
| Routine Restraint | 1.76 | 1.88 | 1.58 | *P*=.27 |
| Compensatory Restraint | 2.09 | 2.16 | 1.98 | *P*=.33 |
| Susceptibility to External Cues | 1.94 | 1.83 | 2.09 | *P*=.55 |
| Emotional Eating | 1.60 | 1.57 | 1.64 | *P*=.72 |

NOTE: P-values estimated by independent t-tests or chi-square analyses. BMI=Body Mass Index; BIS = Barratt Impulsiveness Scale<sup>1</sup>; CESD=The Center for Epidemiological Studies Depression Scale<sup>3</sup>; SHAPS = Snaith-Hamilton Pleasure Scale<sup>6</sup>; PANAS = Positive and Negative Affect Schedule<sup>4</sup>; WREQ=Weight-Related Eating Questionnaire<sup>2</sup>.

**Supplementary Table 2.** The Unadjusted quasi Poisson Regression Analysis showed a significant (P=.01) difference in eating behavior by cluster allocation.

| **term** | **estimate** | **std.error** | **statistic** | ***P* value** |
| --- | --- | --- | --- | --- |
| Intercept | 2.130 | 0.255 | 8.338 | 0.00 |
| Cluster Allocation<br>(baseline: sign-trackers) | 0.868 | 0.324 | 2.680 | 0.01 |

**Supplementary Table 3.** The quasi-Poisson Regression Analysis showed the presence of significant (P=.024) differences in eating behavior by cluster allocation after adjusting for age, gender, BMI, and pre-experiment lever of appetite.

| **term** | **estimate** | **std.error** | **statistic** | ***P* value** |
| --- | --- | --- | --- | --- |
| Intercept | 3.417 | 0.819 | 4.170 | 0.000 |
| Cluster Allocation<br>(baseline: sign-trackers) | 0.802 | 0.343 | 2.337 | 0.024 |
| Age | -0.031 | 0.014 | -2.156 | 0.037 |
| Gender | -0.283 | 0.346 | -0.819 | 0.417 |
| BMI | 0.008 | 0.018 | 0.460 | 0.648 |
| Pre-experiment<br>Level of Appetite | -0.003 | 0.005 | -0.594 | 0.555 |

**Supplementary Figure 1.**
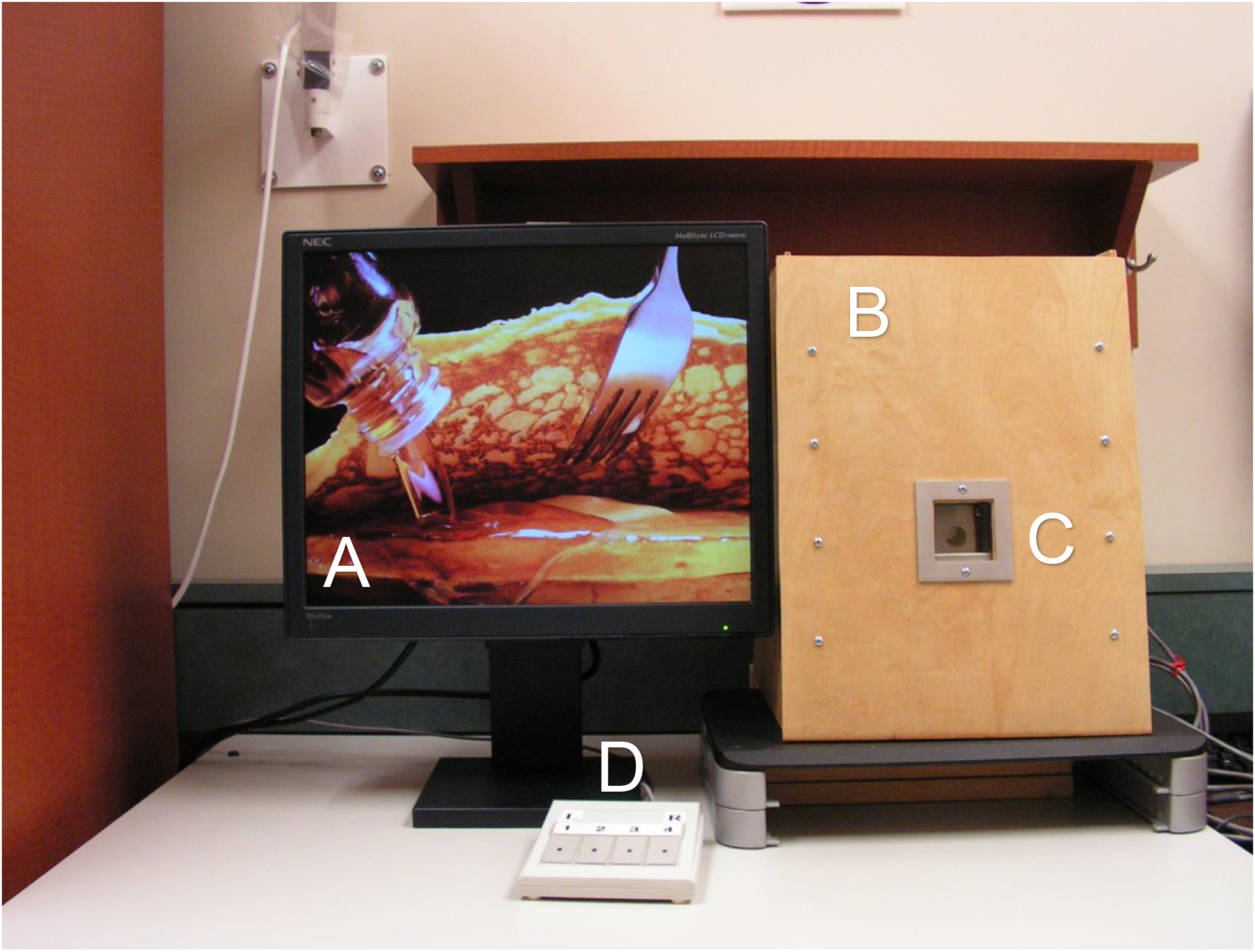
During the cued food delivery task, images appear on the screen (A). The M&M’s^®^ dispenser (behind box B) delivers one candy at a time in the receptacle (C). Participants can either eat or discard the candy. Using the button box (D), the participant can resume the task after eating or discarding the candy.

**Supplementary Figure 2.**
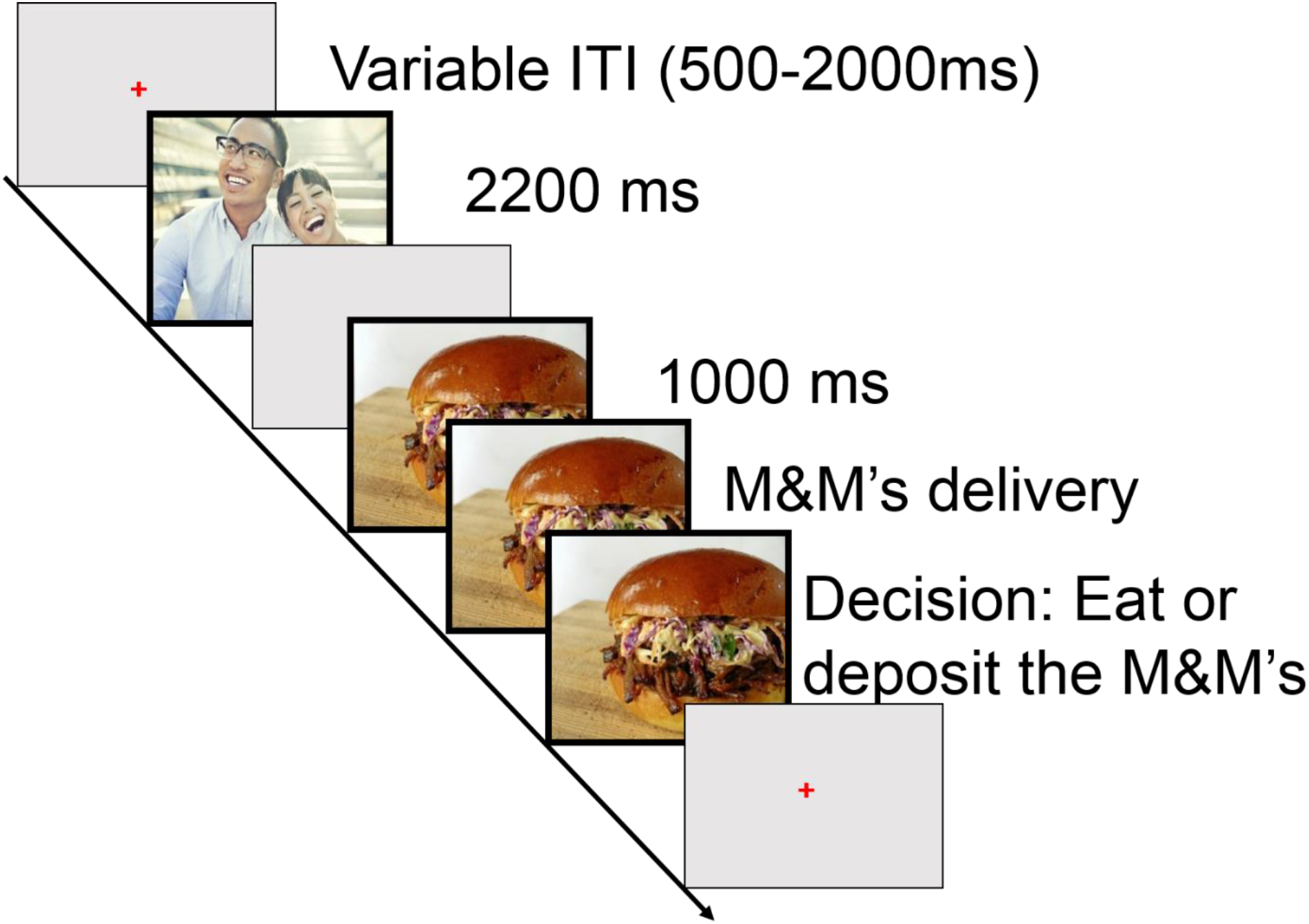
A schematic example of the experimental time course. Note that food predictive cues were presented until the participant decided whether to consume or discard the M&M^®^ candy.

**Supplementary Figure 3.**
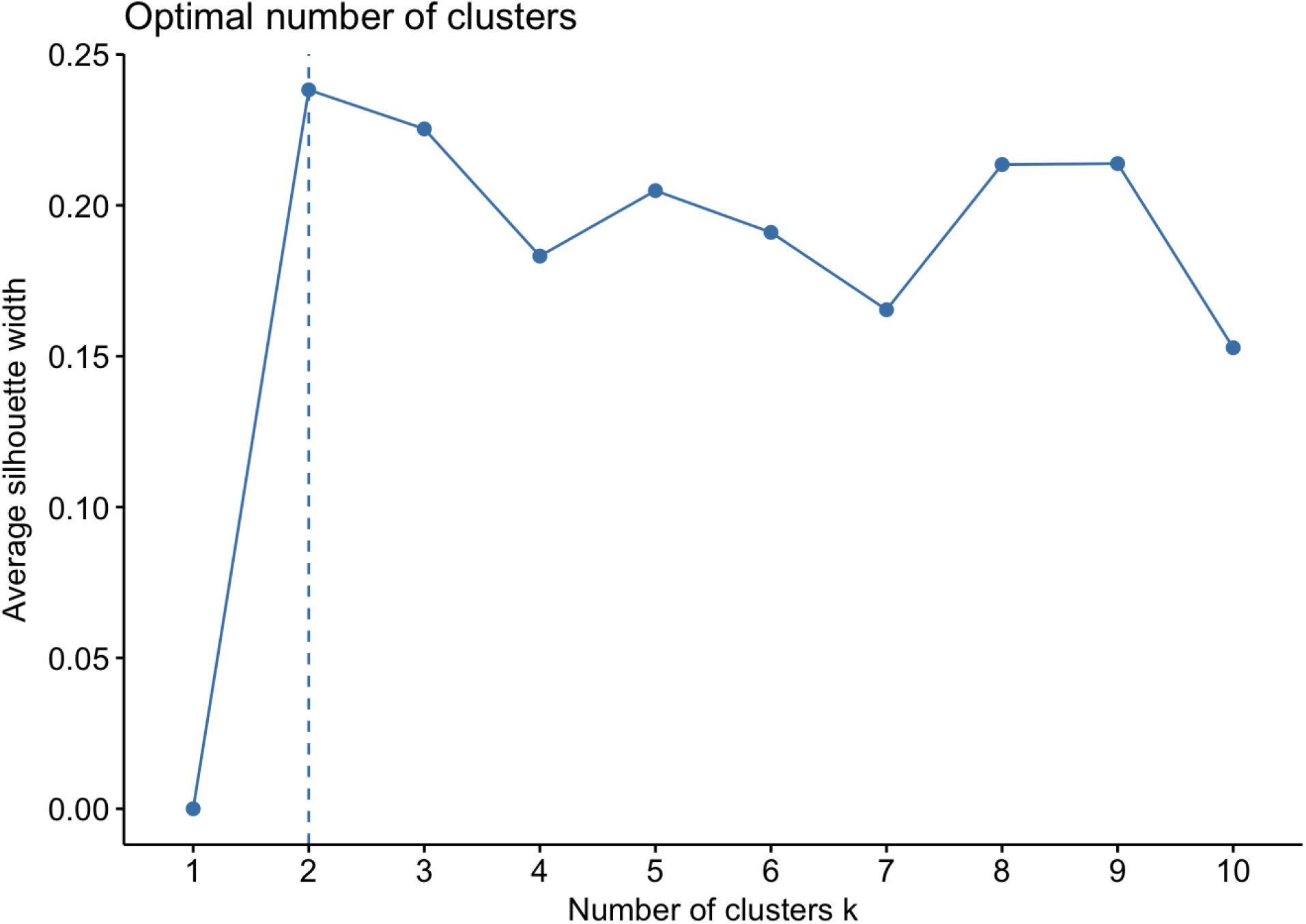
The silhouette method indicated that two was the optimal number of clusters identified by the *k*-means cluster analysis using the 8 LPP values.

**Supplementary Figure 4.**
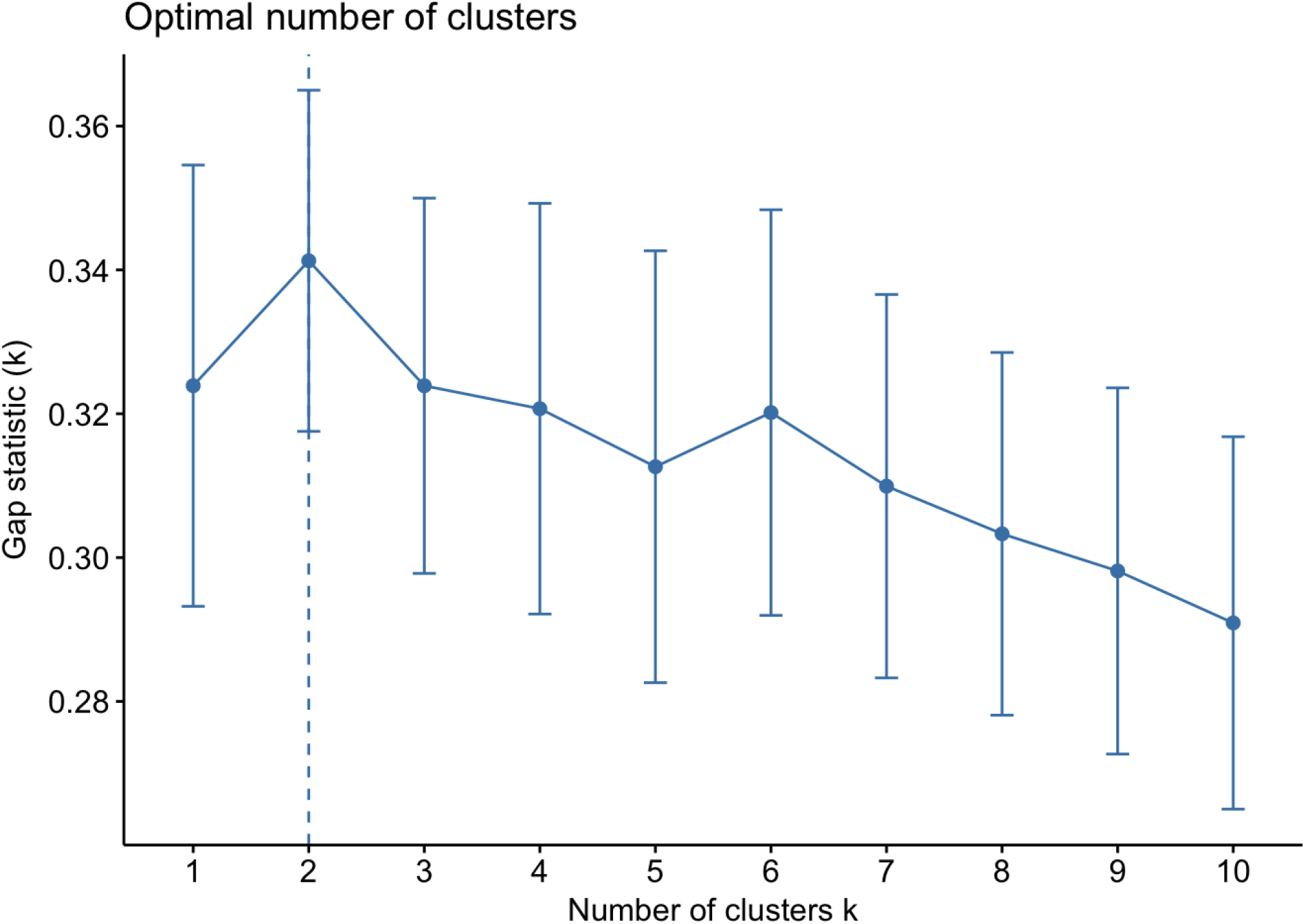
The gap statistic method indicated that two was the optimal number of clusters identified by the *k*-means cluster analysis using the 8 LPP values.

